# Compensating for a shifting world: evolving reference frames of visual and auditory signals across three multimodal brain areas

**DOI:** 10.1101/669333

**Authors:** Valeria C. Caruso, Daniel S. Pages, Marc A. Sommer, Jennifer M. Groh

## Abstract

Stimulus locations are detected differently by different sensory systems, but ultimately they yield similar percepts and behavioral responses. How the brain transcends initial differences to compute similar codes is unclear. We quantitatively compared the reference frames of two sensory modalities, vision and audition, across three interconnected brain areas involved in generating saccades, namely the frontal eye fields (FEF), lateral and medial parietal cortex (M/LIP), and superior colliculus (SC). We recorded from single neurons in head-restrained monkeys performing auditory- and visually-guided saccades from variable initial fixation locations, and evaluated whether their receptive fields were better described as eye-centered, head-centered, or hybrid (i.e. not anchored uniquely to head- or eye-orientation). We found a progression of reference frames across areas and across time, with considerable hybrid-ness and persistent differences between modalities during most epochs/brain regions. For both modalities, the SC was more eye-centered than the FEF, which in turn was more eye-centered than the predominantly hybrid M/LIP. In all three areas and temporal epochs from stimulus onset to movement, visual signals were more eye-centered than auditory signals. In the SC and FEF, auditory signals became more eye-centered at the time of the saccade than they were initially after stimulus onset, but only in the SC at the time of the saccade did the auditory signals become *predominantly* eye-centered. The results indicate that visual and auditory signals both undergo transformations, ultimately reaching the same final reference frame but via different dynamics across brain regions and time.

**New and Noteworthy:** Models for visual-auditory integration posit that visual signals are eye-centered throughout the brain, while auditory signals are converted from head-centered to eye-centered coordinates. We show instead that both modalities largely employ hybrid reference frames: neither fully head-nor eye-centered. Across three hubs of the oculomotor network (intraparietal cortex, frontal eye field and superior colliculus) visual and auditory signals evolve from hybrid to a common eye-centered format via different dynamics across brain areas and time.

## INTRODUCTION

The locations of objects we see or hear are determined differently by our visual and auditory systems. Yet we perceive space as unified, and our actions do not routinely depend on whether they are elicited by light or sound. This suggests that sensory modality-dependent signals ultimately become similar in the lead-up to action. This study addresses how, where, and when this happens in the brain. In particular, we focus on the neural representations that permit saccadic eye movements to both visual and auditory targets.

Differences between visual and auditory localization originate at the sensory organs. Light from different directions excites different retinal receptors, producing an eye-centered map of space that is passed along to thalamic and cortical visual areas. By contrast, sound source localization involves computing differences in sound intensity and arrival time between the two ears, and evaluating spectral cues arising from the filtering action of the outer ear. These physical cues are head-centered and independent of eye movements.

How signals are transformed from one reference frame to another is a comparative question. Many brain areas, including those in the primate oculomotor system, exhibit signals that are not clearly well characterized by a single reference frame either at the neuron or population levels (e.g. Andersen and Mountcastle 1983; Caruso et al. 2018a; Chang and Snyder 2010; Chen et al. 2013; DeSouza et al. 2000; Duhamel et al. 1997; Jay and Sparks 1987b; Lee and Groh 2012; Monteon et al. 2013; Mullette-Gillman et al. 2005, 2009; Russo and Bruce 1994; Sajad et al. 2016; Schlack et al. 2005; Stricanne et al. 1996; Van Opstal et al. 1995; Zirnsak et al. 2014). This makes the signal flow across different brain areas unclear, and requires a quantitative comparison across studies using the same experimental methods.

Here, we compared the reference frames of visual and auditory signals during a sensory guided saccade task across three main hubs in the network controlling saccade target selection and execution. We examined new experimental data from frontal eye fields (FEF), as well as previously presented findings from the main input and output regions to the FEF (figure 1A, Robinson, 1968): the medial and lateral intraparietal cortices (M/LIP^1^) and superior colliculus (SC) (Caruso et al. 2018a; Lee and Groh 2012; Mullette-Gillman et al. 2005, 2009). We found that hybrid signals (i.e. not purely head- or eye-centered) were common in both modalities and across all three brain areas. Visual and auditory signals differed from each other during most temporal epochs, but reached a similar eye-centered format via different dynamics through M/LIP to FEF to SC.

**Figure 1:**
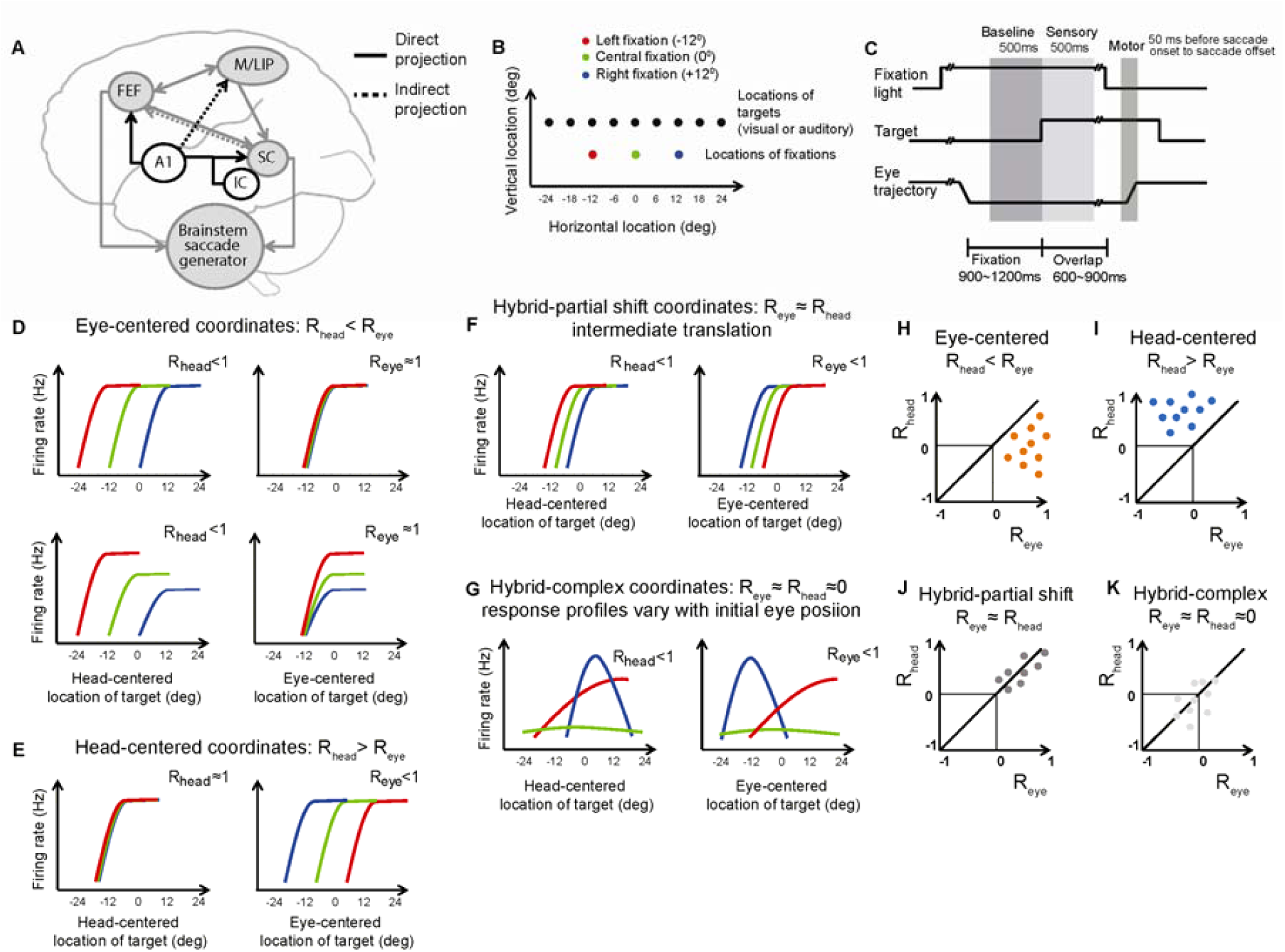
Rationale of the study: the brain areas, stimuli, task and quantification of reference frame. **(A)** Anatomical connections and auditory inputs of M/LIP, FEF and SC. FEF and M/LIP receive auditory inputs from the primary auditory cortex (A1). The SC receives auditory inputs from the inferior colliculus (IC), auditory cortex, M/LIP, and FEF. **(B)** Locations of stimuli and initial fixations. All visual stimuli were green lights: the different colors of the fixation lights in the schematics serve to distinguish tuning curves constructed from different initial fixations in the following graphs. **(C)** Task: Each trial starts with the appearance of a light which the monkey is required to fixate. After a variable delay of 900 to 1200 ms, a visual or auditory target is presented. After a second variable delay of 600 to 900 ms, the fixation light disappears and the monkey reports the location of the target by saccading to it. **(D-G)** Schematics of the relative alignment of the tuning curves from three initial fixation positions plotted in head-and eye-centered coordinates. The strength of the tuning curve alignment in eye- and head-centered coordinates is quantified with the indices R_eye_ and R_head_, which reflect the average correlation between tuning curves. **(D**) **Eye-centered coordinates**. The three tuning curves align well in eye-centered coordinates (R_eye_≈1, right panels), and are separated by the distance between the initial eye positions in head-centered coordinates (R_head_≈0, left-panels). Gain differences across tuning curve do not contribute to the metric chosen to quantify their alignment (lower panels). **(E) Head-centered coordinates**. The pattern is the opposite of (D). **(F) Hybrid-partial shift coordinates**. The three tuning curves are not perfectly aligned in either head- or eye-centered coordinates, but are separated by less than the distance between the fixation locations in both coordinate systems. **(G) Hybrid-complex coordinates**. Both the shape and the alignment vary with the initial eye location. **(H-K) Assessment of reference frame trough the statistical comparison of R**_**eye**_ **vs. R**_**head**_ **for each cell**. The coordinates systems are classified as **(H)** eye-centered if R_eye_>R_head_ (orange dots), **(I)** head-centered if R_head_>R_eye_ (blue), **(J)** hybrid-partial shift if R_eye_≈R_head_≠0 (dark grey), **(K)** hybrid complex if R_eye_≈R_head_≈0 (light gray). The quantitative comparison was carried out via a bootstrap analysis (see Methods).

**Figure 2:**
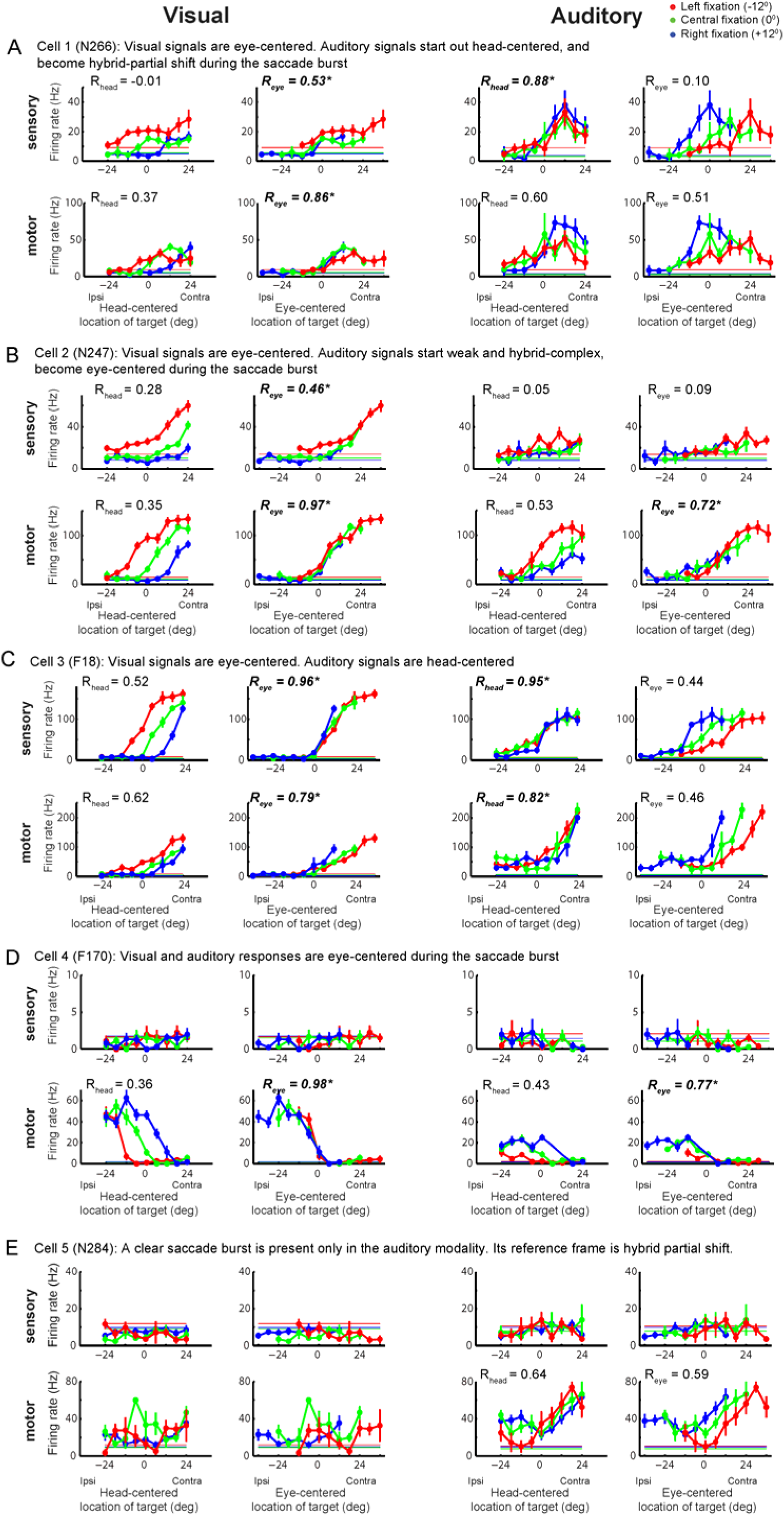
Examples of responses in the FEF. Each panel shows the three tuning curves from left fixation (in red), central fixation (in green) and right fixation (in blue) for an example cell during the sensory and motor period and in the visual and auditory modalities. The tuning curves are plotted in head-centered coordinates (left) and eye-centered coordinates (right). Note that sensory and motor panels have different scales. The horizontal lines indicate the baseline firing rate. The reference frame indices R_head_ and R_eye_ are indicated. See main text for a full description.

## MATERIALS AND METHODS

The behavioral task, neural recording procedures, and analysis methods were identical across brain areas (Caruso et al., 2016, 2018a) (Mullette-Gillman et al., 2005, 2009)(Lee and Groh, 2012). All experimental procedures conformed to NIH guidelines (2011) and were approved by the Institutional Animal Care and Use Committee of Duke University or Dartmouth College. Portions of the data have been previously presented (Mullette-Gillman et al. 2005, 2009; Lee and Groh 2012; Caruso et al. 2018a).

## Experimental Design

Six monkeys (two per each brain area) performed saccades to visual or auditory targets from a range of initial eye locations. Each trial started with the onset of a visual fixation light. On a correct trial, the monkey initiated fixation within 3000 ms. During fixation, after a delay of 900 to 1200 ms, a visual or auditory target turned on in one of nine positions (figure 1B, see below). After a second delay of 600 to 900 ms, the fixation light turned off, while the target stimulus stayed on. The monkey was required to saccade towards the target within 500ms from fixation offset. To prevent exploratory saccades, the money had to keep fixation on the target for 200 to 500 ms. Then the target turned off and the monkey was rewarded with a few drops of diluted juice or water.

The delay (600-900 ms) between target onset and fixation stimulus offset permitted the dissociation of sensory-related activity (i.e. time-locked to stimulus onset) from motor-related activity (i.e. time-locked to the saccade). The targets were placed in front of the monkeys at eye level (0° elevation) and at −24°, −18°, −12°, −6°, 0°, +6°, +12°, +18°, +24° relative to the head along the horizontal direction (figure 1B). The three initial fixation lights were located at −12°, 0°, +12° along the horizontal direction. The elevation of the fixation ranged from −12° to −4° below eye level and from +6° to +14° above eye level. For each isolated neuron, the fixation elevation was chosen as follows. The monkey performed a few prescreening trials to two targets located at [-12°,0°] or [0°,+12°], and starting from two fixation locations, one above and one below eye-level (symmetrical, e.g. [0°,-6°], [0°,+6°]). We chose the fixation elevation (above or below) that was accompanied by qualitatively stronger motor bursts.

The auditory stimuli consisted of white noise bursts (band-passed between 500 Hz and 18 kHz) produced by Cambridge SoundWorks MC50 speakers at 55 dB spl. The visual stimuli consisted of small green spots (0.55 minutes of arc, luminance of 26.4 cd/m2) produced by light emitting diodes (LEDs).

## Recordings

All monkeys (adult rhesus macaques) were implanted with a head holder to restrain the head, a recording cylinder to access the area of interest (M/LIP, FEF or SC) and a scleral search coil to track eye movements. In one of the monkeys, a video eye-tracking system (EyeLink 1000; SR Research, Ontario, Canada) substituted the search coil in a minority of recording sessions (62 over 171, Caruso et al., 2016). The recording cylinders were placed over the left and right M/LIP (monkeys B and C), over the left or the right FEF (monkeys F and N) and over the left and right SC (monkeys W and P). Locations of the recording chambers were confirmed with MRI scans and by assessment of functional properties of the recorded cells (Caruso et al., 2016; Mullette-Gillman et al., 2005, 2009). The behavioral paradigm, and the recordings of eye gaze and of single cell spiking activity were directed by the Beethoven program (Ryklin Software). The trajectory of eye gaze was sampled at 500 Hz. Extracellular activity of single neurons was acquired with tungsten micro-electrodes (FHC, 0.7 to 2.5 MOhm at 1 kHz). A hydraulic pulse microdrive (Narishige MO-95) controlled the electrodes position. A Plexon system (Sort Client software, Plexon) controlled the recordings of single neurons spiking activity.

## Datasets

The basic auditory properties and the visual reference frame of the FEF data set have been previously described (Caruso et al., 2016, 2018a). The analysis of the auditory reference frame is novel. In total, we analyzed 324 single cells from the left and right FEF of two monkeys (monkey F, male, and monkey P, female).

The data from M/LIP and SC have been previously described (Mullette-Gillman et al., 2005, 2009) and are re-analyzed here to permit comparison between all three brain regions. In total, we re-analyzed 179 single cells from the left and right SC of two monkeys (monkey W, male, and monkey P, female), and 275 single cells from the left and right M/LIP of two other monkeys (monkey B, male, and monkey C, female).

### Statistical Analysis

All analyses were carried out with custom-made routines in Matlab (the MathWorks Inc.). Only correct trials were considered. The physical positions of the targets were used for all analyses. Although saccades to auditory targets tend to be more variable than those to visual targets (e.g. Caruso et al. 2016), using saccade endpoints instead of actual target locations did not change the general pattern of results reported here.

For all brain areas, we defined a **baseline**, 500 ms of fixation before target onset, and two response periods. The **sensory period** comprised the 500 ms following target onset (thus including both transient and sustained responses). The pattern of results did not change when the sensory period was shortened to 200 ms (as also indicated by the time course analysis in figures 3G-H and 7). The **motor period** varied according to the temporal profile of the saccade burst: from 150 ms (M/LIP) or 50 ms (FEF) or 20 ms (SC) before saccade onset to saccade offset. Saccade onset and offset were measured as the time in which the instantaneous speed (sampled at 2 ms resolution) of the eye movement crossed a threshold of 25°/s. We also analyzed the entire interval between target onset and saccade using sliding windows of 100 ms (with 50 ms steps), as shown in figures 3G-H and 7.

**Figure 3:**
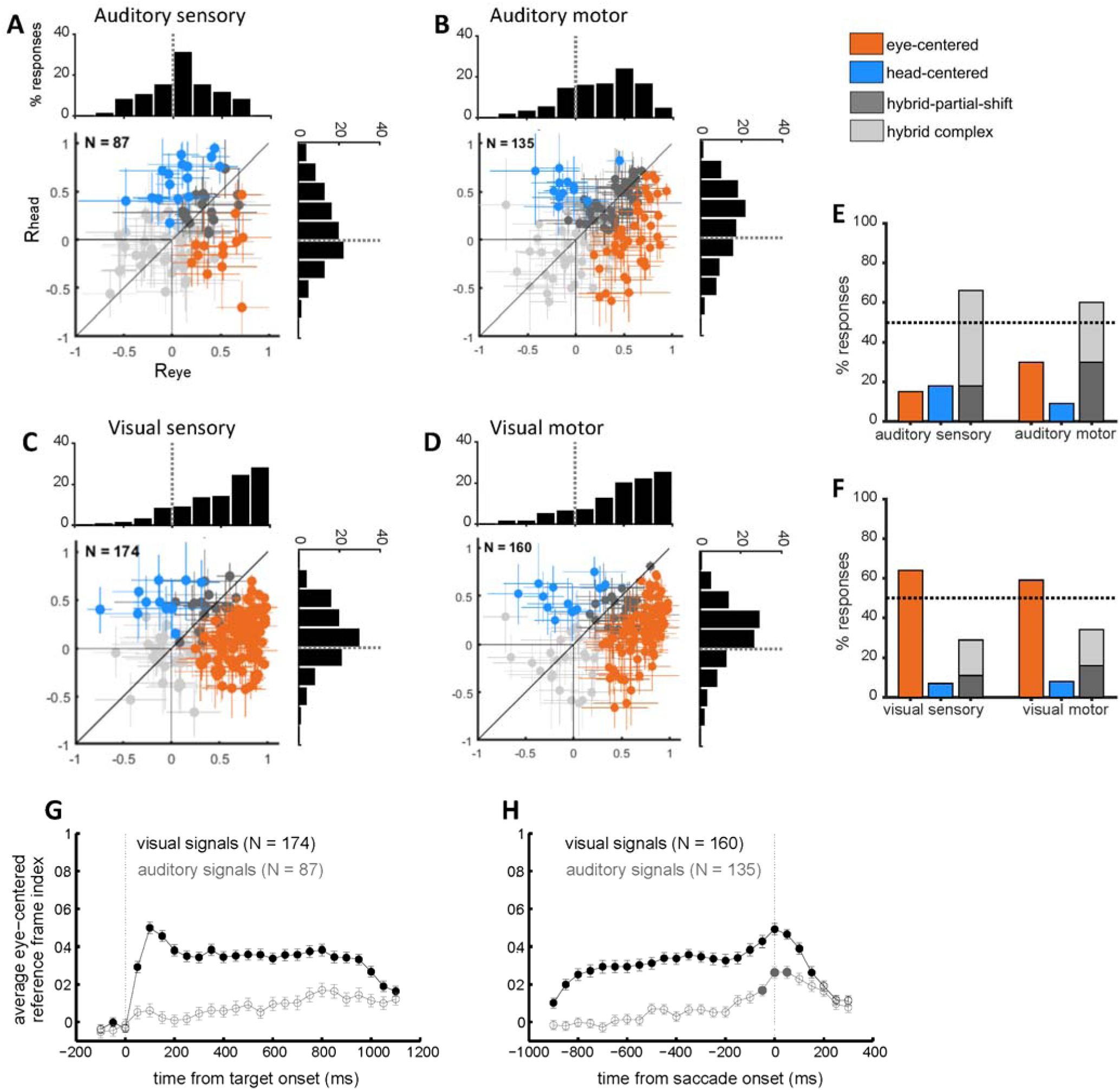
FEF single cell reference frames for visual and auditory targets. **(A-D)** The reference frame indices in head-centered and eye-centered coordinates (R_head_ and R_eye_) are plotted for each individual neuron’s response to visual and auditory targets, in the sensory and motor periods. **(A)** auditory sensory, **(B)** auditory motor, **(C)** visual sensory, **(D)** visual motor. Responses are classified and color-coded as eye-centered, head–centered, hybrid-partial shift and hybrid-complex, based on the statistical comparison between their R_eye_ and R_head_. The histogram inserts show the distribution of R_eye_ (horizontal histogram) and R_head_ (vertical histogram) across the population of cells. **(E-F)** Percentage of auditory (E) and visual (F) responses classified as eye-centered, head–centered, hybrid-partial shift and hybrid-complex, during the sensory and motor periods. Color code as in A. **(G-H)** Time course of the average reference frame index (mean ± SE) in eye-centered coordinates (R_eye_) for the visual and auditory populations, aligned to target onset **(G)** and saccade onset **(H)**. The R_eye_ are calculated in bins of 100 ms, sliding with a step of 50 ms. Filled circles indicate bins in which R_eye_ was significantly greater than R_head_ (as assessed with a t-test, p-value <0.05). Visual data were previously presented in Caruso et al. 2018a.

### Spatial selectivity

The reference frame can only be assessed if the neural response is modulated by target locations. Thus, for each modality and response period, we assessed **responsiveness** with a two-tailed t-test between baseline and response periods, and **spatial selectivity** with 2 two-way ANOVAs measuring the effect of target and fixation location in head- and in eye-centered coordinates on the responses. The inclusion criteria was either a significant main effect for target location or a significant interaction between target and fixation locations, at an uncorrected level of 0.05 for either ANOVA. (Mullette-Gillman et al. 2005, Lee and Groh 2012; Caruso et al. 2018a).

### Reference frame analysis

To determine the reference frame of visual and auditory signals, we quantified the relative alignment between the tuning curved computed in eye-centered or head-centered coordinates for individual neurons. To be specific, for each cell, modality and response period, we computed the three response tuning curves for the three initial fixations twice: once with target locations defined in head-centered coordinates, and once with target locations defined in eye-centered coordinates. For each triplet of tuning curves, we quantified the relative alignment by defining a **reference frame index** according to equation 1. (Mullette-Gillman et al. 2005).

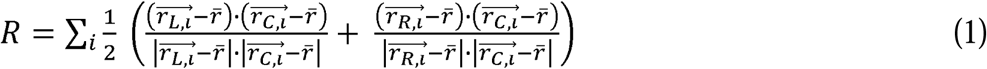

In equation 1, r_L,i_, r_C,i_, and r_R,i_ are the vectors of the time-average responses of the neuron to a target at location *i* and fixation on left (L), right (R), or center (C), and *r* □is the average response across all fixation and target locations. The time windows for the computation of R were the sensory and motor periods as defined above, for the analyses reported in figures 2, 3A-F, 4 and 5, and a 100 ms long sliding window for figures 3G-H and 7. The values of R vary from −1 to 1, with 1 indicating perfect alignment, 0 indicating no alignment and −1 indicating perfect negative correlation.

We statistically compared R_head_ and R_eye_ with a bootstrap analysis (1000 iterations of 80% of data for each target/fixation combination). This let us classify reference frames as:

**1 eye-centered**: if R_eye_ was statistically greater than R_head_

**2 head-centered**: if R_head_ was statistically greater than R_eye_

**3 hybrid-partial shift**: if R_eye_ and R_head_ were not statistically different from each other, but they were different from zero

**4 hybrid-complex**: R_eye_ and R_head_ were not statistically different from zero.

Note that the index R is similar to a correlation coefficient: it measures the relative translation of the three tuning curves, and is invariant to changes in gain as long as the recording captures a part of the receptive field (figure 1D). This was ensured here by including only spatially selective responses as noted above: flat responses that do not vary with target location were not considered.

Only 5 target locations (−12°, −6°, 0°, +6°, and +12°) were included in the analysis, as these were present for all fixation positions in both head- and eye-centered frames of reference. This is a critical aspect of any reference frame comparison because unbalanced ranges of sampling can bias the results in favor of the reference frame sampled with the broader range. In the current study, this bias would have favored an eye-centered reference frame.

Note that any vertical shifts in the receptive field position that might accompany shifts in horizontal fixation were not assessed in this study. If sizeable, such shifts would likely result in the neuron under study being classified as hybrid-complex or hybrid-partial shift.

## Data availability

The data and computer code that support the findings of this study are available from the corresponding authors upon reasonable request.

## RESULTS

### 1. Overview

We quantitatively evaluated the reference frames of visual and auditory targets represented in three brain areas (M/LIP, FEF and SC) at target onset and during a saccade execution. We integrated new data (auditory representations in FEF) with previously reported data (Mullette-Gillman et al. 2005, 2009; Lee and Groh 2012; Caruso et al. 2016; Caruso et al. 2018a).

The task and recording techniques were the same across all brain areas: single unit recordings in 2 monkeys performing saccades to visual or auditory targets at variable horizontal locations and from different initial eye positions (figure 1B-C).

Figure 1D-K shows a summary of possible reference frames exemplified by three hypothetical tuning curves (color coded by the initial gaze direction and plotted in head-centered or eye-centered coordinates). Eye-centered tuning curves align well in eye-centered coordinates (figure 1D), while head-centered tuning curves align in head-centered coordinates (figure 1E). We show two possible ways in which cells deviated from this schema. Namely, we define a *hybrid-partial shift* frame, in which tuning curves with similar shapes are not perfectly aligned in eye- or head-centered coordinates (figure 1F), and a *hybrid-complex* frame, in which tuning curves vary in shape and location when the initial fixation changes (figure 1G). This classification was performed by measuring and comparing the relative alignment of each cell’s three tuning curves in head- and eye-centered coordinates with the reference frame indices, R_head_ and R_eye_, as shown in figures 1H-K. In eye-centered representations R_eye_ is statistically greater than R_head_; in head-centered representations the opposite relation holds (R_head_ > R_eye_); in hybrid-partial shift representations the two indices are both statistically greater than zero but not different from each other (R_eye_ ≈ R_head_ ≠ 0); and in hybrid-complex representations the two indices are not statistically different from zero (R_eye_ ≈ R_head_ ≈ 0).

We included single neuron responses only if we were able to sample at least part of the receptive field during the recordings. Neurons were excluded if they did not exhibit responsiveness (t-test comparing responses to all targets during relevant response period to baseline) and spatial selectivity to target location (ANOVAs; see Methods).

We first present the new data concerning the auditory reference frame in FEF, then turn to how visual and auditory signals change across brain areas.

### 2. FEF auditory response patterns are predominantly hybrid

Individually, neurons in the FEF showed considerable heterogeneity of reference frame. Figure 2 shows visual and auditory response profiles at target onset and at saccade execution for 5 example cells. In figure 2A-C, visual responses are eye-centered, while auditory responses vary across cells and response period. In figure 2A, auditory signals are head-centered at target onset (compare R_head_ and R_eye_: 0.88 vs. 0.1) and become hybrid-partial shift during the saccade (R_head_=0.60 vs. R_eye_=0.51). In figure 2B, auditory responses are hybrid-complex at target onset and become eye-centered during the saccade. Finally, in figure 2C, auditory signals are head-centered in both periods. Figure 2D-E shows two cells that did not respond to visual and auditory targets, but fired during saccades. In figure 2D, such motor activity was classified as eye-centered in both modalities. In figure 2E, the auditory motor activity was classified as hybrid-partial shift, while the visual motor activity was not classified as it did not pass the tuning criteria (the response was higher than baseline but did not vary with target location, see Methods).

While individual cells could be found anywhere along the spectrum from head-centered to eye-centered (figure 2, figure 3A-F), the majority of auditory responses in FEF were hybrid, at target onset as well as during saccade execution. When the R_head_ and R_eye_ values for individual neurons are plotted against each other (figure 3A-B), the 95% confidence intervals of those values overlap, indicating that auditory responses from different initial eye positions are not better aligned in eye-centered or head-centered coordinates (gray cross hairs).

Treating reference frame as a category, fewer than 20% of FEF auditory sensory responses were cleanly classified as head-centered (18%) or eye-centered (15%) (orange and blue points in figure 3A-B, orange and blue bars in figure 3E). At saccade execution, the proportion of eye-centered auditory responses doubled to 30%, while head-centered responses decreased to 9%. Thus, in the auditory modality, there was a limited shift in time towards eye-centered representations during saccade execution. However, auditory eye-centered responses remained modest in prevalence compared to auditory hybrid responses (stably around 60% throughout the task, figure 3E), and to visual eye-centered responses (also around 60% throughout the task, figure 3F). The difference between auditory and visual motor reference frames cannot be attributed to a difference in the strength of visual and auditory population responses, as these had a similar average magnitude (Caruso et al., 2016).

The shift in auditory reference frame and the differences between visual and auditory coding were also evident in the distribution of values of the R_eye_ and R_head_ indices (see histograms in figure 3A-D). For each condition, we compared the population-average magnitude of R_eye_ and R_head_ with a t-test at a Bonferroni-adjusted alpha level of 0.0125 (0.05/4). For auditory signals, R_eye_ was statistically greater than R_head_ in the motor period (t-test, p<0.10^−3^), but not in the sensory period (t-test, p=0.8), while for visual signals R_eye_ was consistently larger than R_head_ (t-test, p<0.10^−3^).

This finding held for finer time scales as well: figures 3G and 3H plot the average population values of R_eye_ and R_head_ across time (100 ms bins). Visual responses were predominantly, though not fully, eye-centered from target onset to saccade completion (Caruso et al., 2018a), while auditory responses only slightly shifted towards eye-centeredness at the time of a saccade and did not reach the level of eye-centeredness seen for visual signals. This indicates that the representations of these two modalities remained discrepant at this brain area and temporal stage of neural processing.

### 3. Evolution of coordinates across M/LIP, FEF, and SC

The pattern of classification of auditory reference frames in the FEF appears intermediate to those we previously assessed in the M/LIP and SC with the same experimental paradigm, shown in figures 4 and 5 (Mullette-Gillman et al. 2005, 2009; Lee and Groh 2012). The comparison of figure 3A-F with figures 4 and 5 suggests that both visual and auditory signals undergo a reference frame transformation across areas: signals in M/LIP are less eye-centered/more hybrid than FEF for both modalities, and more eye-centered/less hybrid in the SC than FEF. However, although the final stage of this transformation is represented by eye-centered motor signals in the SC, the evolution across area and time appears different for visual and auditory signals: there is little change in reference frame across time for either modality in M/LIP whereas in the SC visual signals can largely be categorized as eye-centered in both temporal epochs and auditory signals become eye-centered by the time of the saccade.

**Figure 4:**
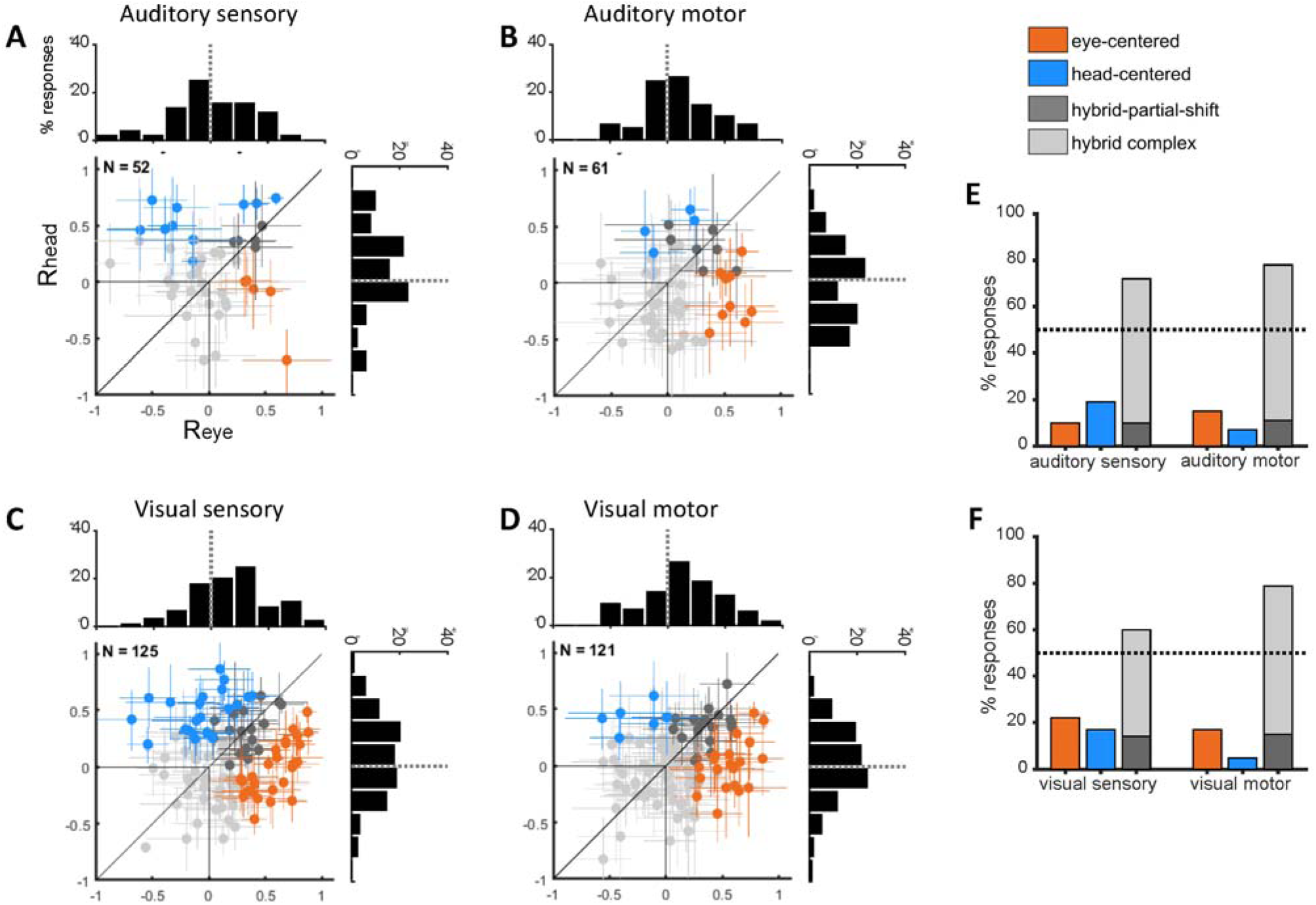
M/LIP single cell reference frames for visual and auditory targets. Same format as figures 3A-F. Data from Mullette-Gillman et al., 2005, 2009.

**Figure 5:**
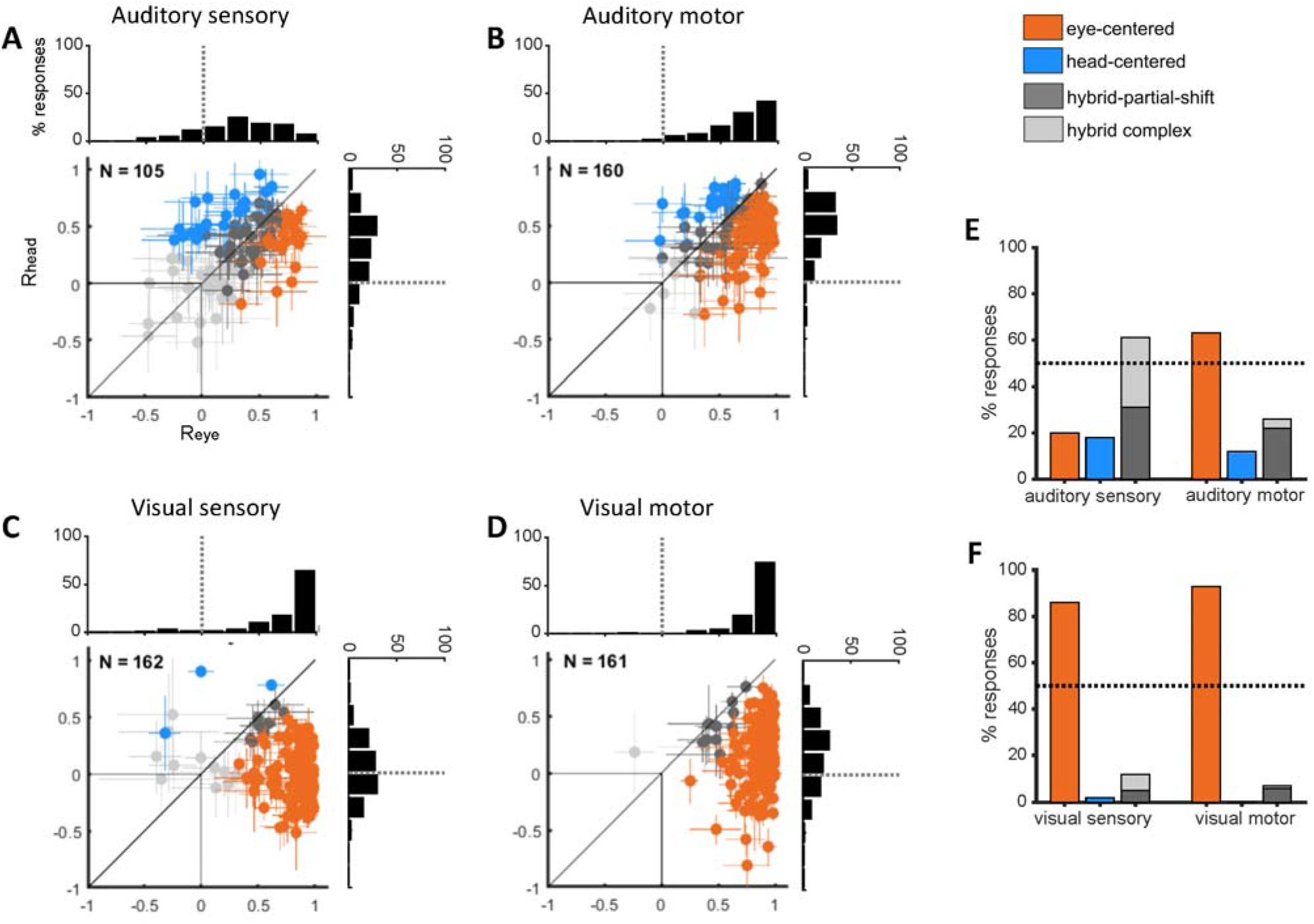
SC single cell reference frames for visual and auditory targets. Same format as figures 3A-F. Data from Lee and Groh, 2012.

To quantify the dynamics of coordinate transformation across modalities, we statistically compared the degree of eye-centeredness (mean R_eye_) by areas (M/LIP, FEF, SC), response period (sensory, motor), and target modality (visual, auditory) with a three-way ANOVA (Figure 6 A-B and Table 1). Neural representations of target location became more eye-centered along the pathway from M/LIP to FEF to SC, but this trend was different across modality (R_eye_ was higher and increased more steeply for visual signals, as indicated by the significant interaction between area and modality) and for sensory vs. motor responses (R_eye_ was higher and increased more steeply for motor responses, as indicated by the significant interaction between area and response period). For all areas, the degree of eye-centeredness of visual signals was fairly stable across sensory and motor periods, while auditory signals became more eye-centered during the motor period (interaction between modality and response period).

**Table 1.** Three-way ANOVA measuring the effect of recording location (M/LIP, FEF, SC), target modality (visual vs. auditory) and response period (target onset vs. saccade) on the average R_eye_. The data were Fisher-transformed for the analysis.

| Effect | d.f. | F | p | $\eta^2$ |
| --- | --- | --- | --- | --- |
| area | 2 | 343.1 | <10-3 | 0.253 |
| modality | 1 | 226.1 | <10-3 | 0.083 |
| response period | 1 | 34.5 | <10-3 | 0.013 |
| area x modality | 2 | 30.8 | <10<sup>-3</sup> | 0.023 |
| area x period | 2 | 18.3 | <10<sup>-3</sup> | 0.013 |
| period x modality | 1 | 19.3 | <10<sup>-3</sup> | 0.007 |
| area x modality x<br>period | 2 | 0.8 | 0.45 | 0.0006 |

**Figure 6.**
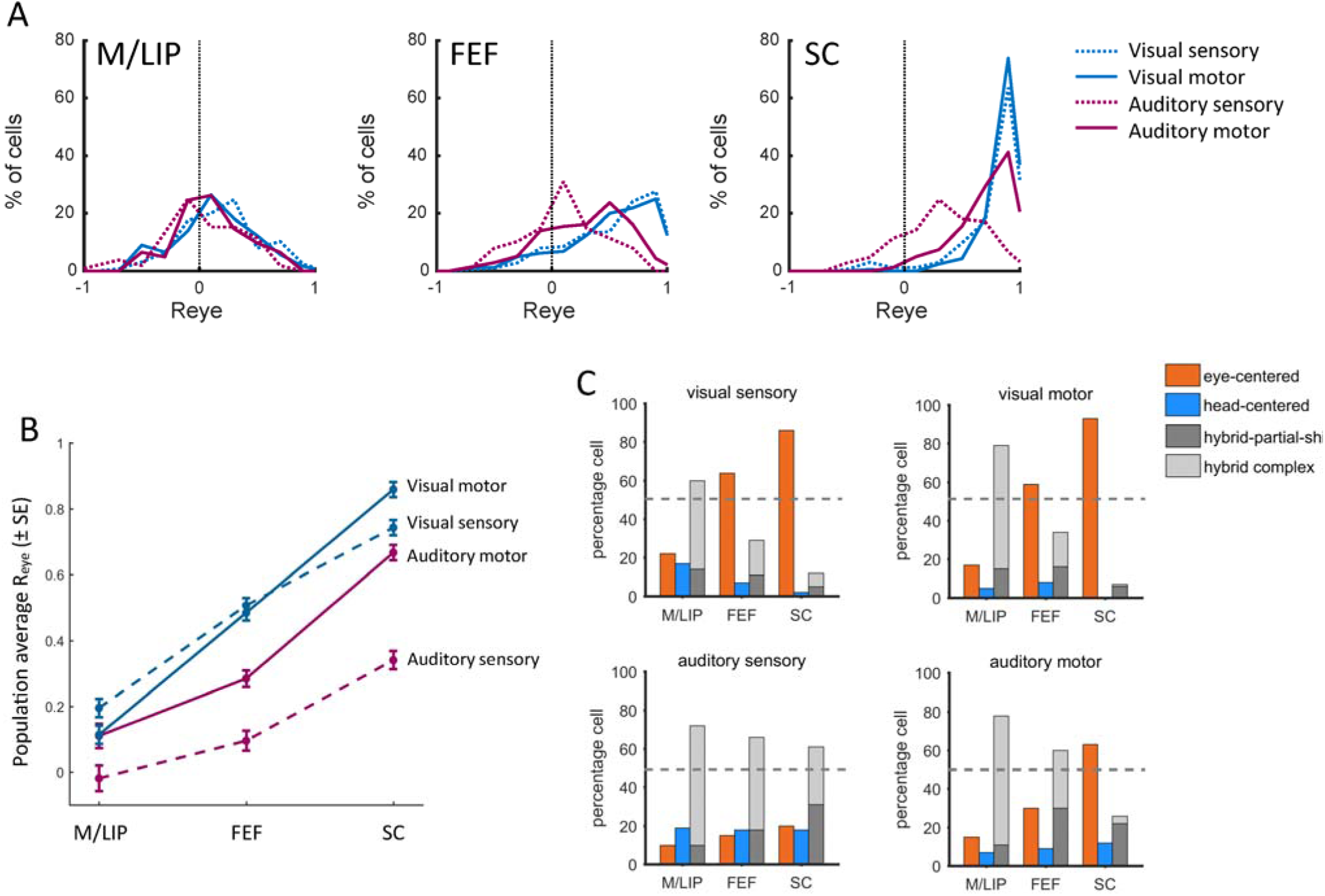
Coordinate transformation across brain areas, modality and response period. (A) Distribution of R_eye_ for visual and auditory signals in the sensory and motor periods in each area (M/LIP, FEF, SC) (B) Average degree of eye-centeredness (mean R_eye_) across neural populations in M/LIP, FEF, SC, for visual and auditory locations, at target onset and at saccade execution.

The classification of individual neuronal responses as eye-centered, head-centered and hybrid across areas, modality and response period, confirmed the different dynamics in the visual and auditory coordinate transformation (figure 6C). The proportion of visual responses classified as eye-centered steadily grew from M/LIP to FEF to SC, while eye-centered auditory responses remained a minority in M/LIP and FEF and increased to over 60% in the SC in the motor period. Furthermore, figure 6C shows that across all areas the proportion of head-centered responses was very low at all times (<20%). Thus, the transformation occurs from hybrid to eye-centered coordinates. Among hybrid responses, the class of hybrid-complex, which captures changes other than partial shifts of the receptive fields, was the biggest category in the M/LIP for both visual and auditory responses, and a sizeable category in FEF and SC during auditory sensory responses. This suggests that the variability of M/LIP neural responses might not be fully understood under the lens of shifting receptive fields.

This general pattern is confirmed by the fine time course of the population R_eye_ across modality and area (figure 7): the degree of eye-centeredness was consistently higher in the SC than in the FEF, which was in turn higher than M/LIP. This was true for both visual (figure 7A) and auditory modalities (figure 7B). However, the SC was the only structure to consistently show an average eye-centeredness that exceeded 0.5, and it did so throughout for visual stimuli, but only at the time of the saccade for auditory stimuli.

**Figure 7:**
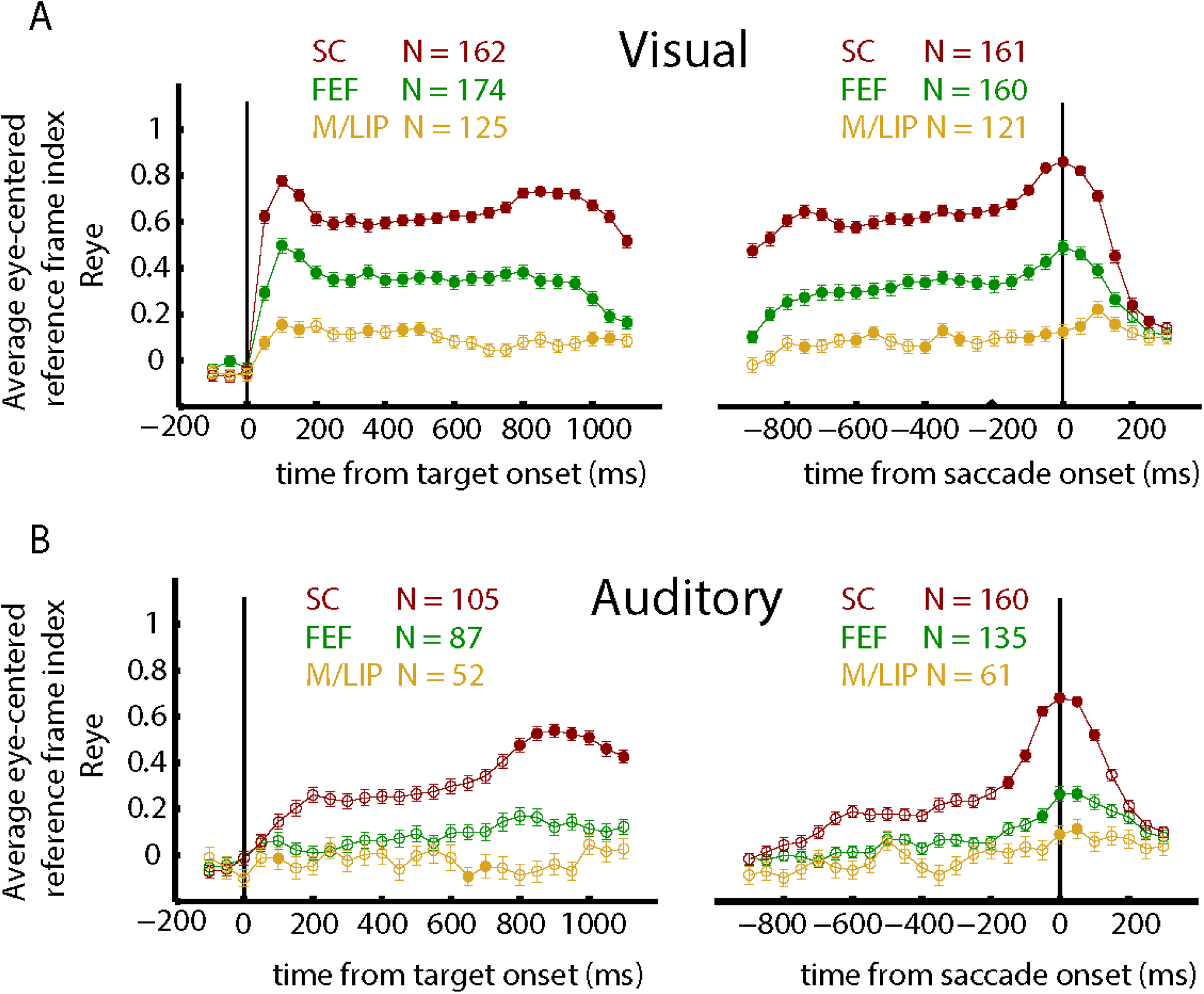
Time course of the eye-centered reference frame indices in M/LIP, FEF, SC. The average eye-centered reference frame index, R_eye_ (mean ± SE) was computed in 100 ms bins (sliding with a step of 50 ms). Filled circles indicate that R_eye_ was statistically larger than R_head_ (see Methods). **(A)** Visual signals, aligned to the target onset (left) and to the saccade onset (right). **(B)** Auditory signals, aligned to the target onset (left) and to the saccade onset (right).

In summary, the coordinate transformation for visual and auditory signals appears to reach a final eye-centered format via different dynamics through M/LIP and FEF to SC: visual signals gradually shift from hybrid to eye-centered, while auditory signals remain hybrid throughout the two cortical levels and only become eye-centered at the SC’s motor command stage.

## DISCUSSION

We compared the reference frames for auditory and visual space in three brain areas (M/LIP, FEF, and SC) during a sensory guided saccade task. At sound onset, auditory signals were not encoded in pure eye-centered or head-centered reference frames, but rather employed hybrid codes in which the neural responses depended on both head-centered and eye-centered locations. During the saccade, auditory responses became more eye-centered in the SC and to a lesser extent in the FEF. The SC was the only region showing a majority of eye-centered auditory motor signals. Hybrid responses were also common in the visual modality, in particular in FEF and M/LIP. They constituted a majority of the response patterns in M/LIP. The proportion of eye-centered visual signals increased in the FEF and SC compared to M/LIP.

To quantify single neuron reference frames, we adopted a correlation approach and measured the degree to which neural signals were anchored to the locations of the eye or the head when these locations were experimentally misaligned. To avoid bias in the estimates of single neurons’ reference frame, we sampled space symmetrically in both coordinate systems, eliminating any “extra” locations that existed in one reference frame but not the other. Without such balanced sampling, the estimate would be erroneously high in favor of the more extensively sampled reference frame. For example, early studies in intraparietal cortex concluded that visual representations were eye-centered with an eye position gain modulation (e.g. Andersen et al. 1985), but a later study suggested that this finding may have been due to inadequate sampling in a head-centered frame of reference (Mullette-Gillman et al. 2009).

The limitations of our approach are related to the possibility that systematic changes in neural responses across populations and task conditions might affect measures of neural signals’ alignment. For example, low response strength, high trial-to-trial variability, and low spatial selectivity would each reduce the correlation measures in both coordinate systems, resulting in more hybrid estimates. However, we did characterize strong eye-centered and head-centered representations in all regions and modalities, indicating that multiple reference frames were active, beyond hybrid signals. Indeed, there is consensus in the literature that eye movements/changes in eye position affect visual and auditory signals in all of the areas we tested here (Andersen and Mountcastle 1983; Andersen et al. 1985, 1987, 1990; Andersen and Zipser 1988; Barash et al. 1991a, b; Batista et al. 1999; Berman et al. 2005, 2007; Cassanello and Ferrera 2007; Cohen and Andersen 2000; Colby et al. 1993, 2005; DeSouza et al. 2000, 2011; Duhamel et al. 1992; Heiser and Colby 2006; Heiser et al. 2005; Jay and Sparks 1984, 1987a,b; Keith et al. 2009; Klier et al. 2001; Lee and Groh 2012; Mullette-Gillman et al. 2005, 2009; Russo and Bruce 1994; Sadeh et al. 2015; Sajad et al. 2015, 2016; Sommer and Wurtz 2006; Stricanne et al. 1996; Van Opstal et al. 1995; Zipser and Andersen 1988; Zirnsak and Moore 2014; Zirnsak et al. 2014).

Differences in how spatial information is coded across sensory systems pose problems for the perceptual integration of the different sensory inputs that arise from a common source, such as the sight and sound of someone speaking. Since the discovery of auditory response functions that shift with changes in eye position (Jay and Sparks 1984), the model for visual-auditory integration has been that visual signals remain in eye-centered coordinates and auditory signals are converted into that same format. Some experimental results, however, cannot be explained under this view. In particular, “pure” reference frames, representations in which the responses are clearly better anchored to one frame of reference over all others, have proved surprisingly rare. In the experiments described here, only the SC contains strongly eye-centered representations, for visual signals during both sensory and motor periods, and for auditory signals only in the motor period. All other observed codes appear to be at least somewhat impure, with individual neurons exhibiting responses that are not well captured by a single reference frame as well as different neurons employing different reference frames across the population. Hybrid, intermediate, impure or otherwise idiosyncratic reference frames have also been observed in numerous other studies (Avillac et al. 2005; Chang and Snyder, 2010; Chen et al., 2013; Mullette-Gillman et al. 2005,2009; Schlack et al. 2005; Stricanne et al. 1996;).

A possible explanation for such hybrid states is that coordinate transformations might be computationally difficult to achieve and hybrid codes represent intermediate steps of a coordinate transformation that unfolds across brain regions and time. Our results lend empirical support to this notion of a gradual, transformative process. However, why such a transformation should unfold across time and brain areas is unclear from a theoretical perspective. Although some models of coordinate transformations involve multiple stages of processing (e.g. Keith et al. 2009; Pouget and Sejnowski 1997; Smith and Crawford 2005; Zipser and Andersen 1988;), there also exist computational models of coordinate transformations in one step (Groh and Sparks 1992). Thus, the computational or evolutionary advantages of such a system remain unclear. Models using time-varying signals such as eye velocity signals to accomplish the coordinate transformation would be a good place to start to explore how this computation unfolds dynamically (e.g. Keith et al. 2009; Droulez and Berthoz, 1991).

Regardless, hybrid codes contain the necessary information to keep track of where targets are with respect to multiple motor effectors, and may therefore facilitate flexible mappings of multiple behaviors or computations on the same stimuli in different contexts (Bernier and Grafton 2010; Crespi et al. 2011). In other words, hybrid representations, at least hybrid partial-shift, are easily read-out codes. As such, hybrid might be the actual representational format for visual and auditory targets in the oculomotor system, rather than an intermediate step in a coordinate transformation. In this scenario, it is plausible that visual and auditory signals become maximally similar in the SC at the moment in which a particular behavioral response, namely a saccade, is executed, as the SC is a key control point in the oculomotor system, not only encoding target location in abstract, but also controlling the time profile and dynamic aspects of saccadic movements (Stanford et al. 1996; van Opstal and Goossens 2008).

The exact mechanism by which auditory signals become predominantly eye-centered in the SC is unknown. Our data do not fully resolve whether a coordinate transformation unfolds locally within the SC, or whether it reflects preferential sampling of eye-centered auditory information through mechanisms of gating or selective read-out. Indeed, the SC gets auditory inputs primarily from FEF and inferior colliculus (IC), both containing a variety of simultaneous auditory representations, from head-centered to eye-centered (see figure 3 and Porter et al. 2006).

Apparent mixtures of different reference frames could also be a consequence of the use of different coding formats across time, with individual neurons switching between different reference frames for different individual epochs of time. Response measures that involve averaging across time and across trials would yield apparently hybrid or intermediate reference frames when in reality, one reference frame might dominate at each instant in time. We recently presented evidence that neurons can exhibit fluctuating activity patterns when confronted with two simultaneous stimuli and suggested that such fluctuations may permit both items to be encoded across time in the neural population (Caruso et al., 2018b). Similar activity fluctuations might permit interleaving of different reference frames across time, with the balance of which reference frame dominates shifting across brain areas and across time. At the moment, this hypothesis is difficult to test because the statistical methods deployed in Caruso et al., 2018b require being able to ascertain the responses to each condition in isolation of the other, but every reference frame exists concurrently. Novel analysis methods, perhaps relying on correlations between the activity patterns of simultaneously recorded neurons, are needed to address this question.

A possible clue that activity fluctuations may be an important factor comes from studies of oscillatory activity. If neural activity fluctuates on a regular cycle and in synchrony with other neurons, the result is oscillations in the field potential. Indeed, studies of the coding of tactile stimuli in saccade tasks provide evidence for different reference frames encoded in different frequency bands, in different brain areas and at different periods of time. In the context of saccade generation, oscillations could allow the selection of one reference frame by adjusting the weights of spatial information received from different populations, in essence flexibly recruiting eye-centered populations (Buchholz et al. 2011, 2013, 2014; Haegens et al. 2011).

In summary, our results suggest that new ways of thinking about how visual and auditory spaces are integrated, beyond simplistic notions of a single pure common reference frame and registration between visual and auditory receptive fields, are needed.

## Acknowledgments

This work was supported by the National Institute of Health: NIH (NIDCD) R01 DC013906; NIH (NIDCD) R01 DC016363, NIH R01 NS50942-05. The authors are grateful to Jessi Cruger, Karen Waterstradt, Christie Holmes and Stephanie Schlebusch for animal care, and to Tom Heil and Eddie Ryklin for technical assistance. The authors have benefitted from thoughtful discussions with Jungah Lee, Kurtis Gruters, Shawn M. Willett, Jeffrey M. Mohl, David Murphy, Bryce Gessell and James Wahlberg.

## Footnotes

1 *Mullette-Gillman et al. 2009* found no difference in reference frame on the basis of recording location in the banks of the IPS, thus we do not distinguish between medial and lateral intraparietal cortex.

